# Hyperpiliation reduces *Pseudomonas aeruginosa* pathogenicity by impeding type III secretion

**DOI:** 10.1101/2021.01.28.428742

**Authors:** Sara L.N. Kilmury, Katherine J. Graham, Ryan P. Lamers, Lesley T. MacNeil, Lori L. Burrows

## Abstract

Type IVa pili (T4aP) are important virulence factors for many bacterial pathogens. Previous studies suggested that the retraction ATPase, PilT, modulates pathogenicity due to its critical role in pilus dynamics and twitching motility. Here we use a *Caenorhabditis elegans* slow killing model to show that hyperpiliation, not loss of pilus retraction, reduces virulence of *Pseudomonas aeruginosa* strains PAK and PA14 by interfering with function of the contact-dependent type III secretion system (T3SS). Hyperactivating point mutations in the *P. aeruginosa* PilSR two-component system that controls transcription of the major pilin gene, *pilA*, increased levels of surface pili to the same extent as deleting *pilT*, without impairing twitching motility. These functionally hyperpiliated PilSR mutants had significant defects in pathogenicity that were rescued by deleting *pilA* or by increasing the length of T3SS needles via deletion of the needle-length regulator, PscP. Hyperpiliated *pilT* deletion or *pilO* point mutants showed similar PilA-dependent impairments in virulence, validating the phenotype. Together, our data support a model where a surfeit of pili prevents effective engagement of contact-dependent virulence factors. These findings suggest that the role of T4aP retraction in virulence should be revised.

**SIGNIFICANCE:** *Pseudomonas aeruginosa* is a major contributor to hospital-acquired infections and particularly problematic due to its intrinsic resistance to many front-line antibiotics. Strategies to combat this and other important pathogens include development of anti-virulence therapeutics. We show that the pathogenicity of *P. aeruginosa* is impaired when the amount of type IVa pili (T4aP) expressed on the cell surface increases, independent of the bacteria’s ability to twitch. We propose that having excess T4aP on the cell surface can physically interfere with productive engagement of the contact-dependent type III secretion toxin delivery system. A better understanding of how T4aP modulate interaction of bacteria with target cells will improve the design of therapeutics targeting components involved in regulation of T4aP expression and function, to reduce the clinical burden of *P. aeruginosa* and other T4aP-expressing bacteria.

## INTRODUCTION

Type IVa pili (T4aP) are versatile surface appendages used for attachment, biofilm formation, DNA uptake, electron transfer, and movement across solid and semi-solid surfaces via twitching motility (1–5). These functions make T4aP important virulence factors for many bacterial pathogens, including the model organism, *Pseudomonas aeruginosa*. T4aP have also been implicated in surface sensing, initiating a signal cascade for virulence factor upregulation (6, 7).

In *P. aeruginosa*, T4aP are composed predominantly of hundreds to thousands of repeating subunits of the major pilin protein, PilA, whose expression is autoregulated by the PilSR two-component system (TCS) (8–11). The lollipop-shaped PilA monomers are anchored in the inner membrane by their N-termini when they are not part of an assembled pilus fibre (12), and are polymerized into a growing pilus by the platform protein PilC and the assembly ATPase PilB, using mechanical energy generated from ATP hydrolysis (13–15). Polymerized pili are then disassembled (retracted) via removal of pilin subunits at the pilus base by PilC, using mechanical energy generated by the retraction ATPase, PilT (16). When *pilT* is absent, pili are assembled but not retracted, resulting in a hyperpiliated strain that is no longer capable of twitching (16, 17).

Multiple studies demonstrated the importance of functional T4aP for *P. aeruginosa* infection of a variety of hosts (8, 18–21). Similar findings have been reported for other T4aP-expressing bacteria, including *Dichelobacter nodosus* (22), *Neisseria meningitidis* (23), *N. gonorrhoeae* (24), *Kingella kingae* (25), *Xanthomonas citri* (26), and *Xylella fastidiosa* (27). Most studies compared the pathogenicity-associated phenotypes of wild-type (WT) strains to isogenic non-piliated mutants. However, the virulence of retraction-deficient *pilT* mutants of *D. nodosus* in sheep (22), *Pantoea ananatis* in onion seedlings (28), *P. aeruginosa* in murine and human infection models (29, 30), *Acidovorax avenae* in seed transmission assays (31), and *N. meningitidis* in mice (32) has been investigated. In each case, *pilT* mutants were less virulent and/or defective in attachment compared to their parent strains. Those studies generally attributed the loss of pathogenicity to loss of pilus retraction and twitching motility, and in most cases, the independent contribution of hyperpiliation to loss of virulence was not considered.

In addition to its role in T4aP retraction, PilT is important for efficient function of the Type III secretion system (T3SS) (33–35). The T3SS machinery forms a rigid needle-like structure that in *P. aeruginosa* allows for injection of the toxic effector proteins ExoS, ExoT, ExoY and/or ExoU directly from the cytoplasm of the bacterium into the cytoplasm of eukaryotic host cells (36). Loss of PilT – or a second putative retraction ATPase in *P. aeruginosa*, PilU – reduced the efficiency of T3S, though the latter had a less significant impact (34). Based on these findings, pilus retraction was proposed to be necessary for establishment of intimate contact with target cells to allow for T3S.

Here, we tested hyperpiliated *P. aeruginosa* mutants with a range of twitching motility capabilities for virulence in a T3SS-dependent *Caenorhabditis elegans* slow killing model. We show that all hyperpiliated mutants tested, regardless of their ability to twitch, were less pathogenic than wild type (WT), and that those virulence defects could be reversed by secondary mutations that abrogated pilus assembly. Based on these results, we propose a model in which hyperpiliation—and not loss of pilus retraction or twitching motility, as previous studies have suggested—reduces the infectivity of *P. aeruginosa*, potentially by impeding the productive engagement of contact-dependent toxin delivery systems including the T3SS.

## RESULTS

### Increased activity of PilSR causes hyperpiliation, without loss of pilus function

The PilSR two-component system controls the transcription of the major pilin gene, *pilA* (9, 10, 37). We showed previously that the system can be constitutively activated using a specific point mutation in the conserved phosphatase motif of PilS, N323A, preventing deactivation of phospho-PilR (11, 38). To test if hyperactivation of PilR also led to increased surface piliation, we generated a point mutant, D54E, which mimics the active, phosphorylated state of the response regulator (39, 40). Both point mutations were introduced at the native *pilSR* loci on the chromosome to ensure WT expression levels. Similar to PilS N323A, the PilR D54E mutant expressed more PilA compared to WT **(Figure 1A).** Analysis of sheared surface pili revealed that both the PilS and PilR point mutants were hyperpiliated **(Figure 1A).** This phenotype resembled that of mutants lacking the retraction ATPase, *pilT* (16), or bearing an M92K mutation in the alignment subcomplex protein, PilO (41). Surface piliation was lost in all strains when the pilin gene, *pilA*, or the gene encoding the assembly ATPase, *pilB*, was deleted. In contrast to the retraction-deficient *pilT* mutant that fails to twitch, or the PilO M92K mutant which has reduced twitching, the PilS and PilR point mutants retain WT levels of twitching, suggesting that their pili are fully functional **(Figure 1C).** This set of hyperpiliated mutants, representing a range of pilus functionality, was used in further studies.

**Figure 1.**
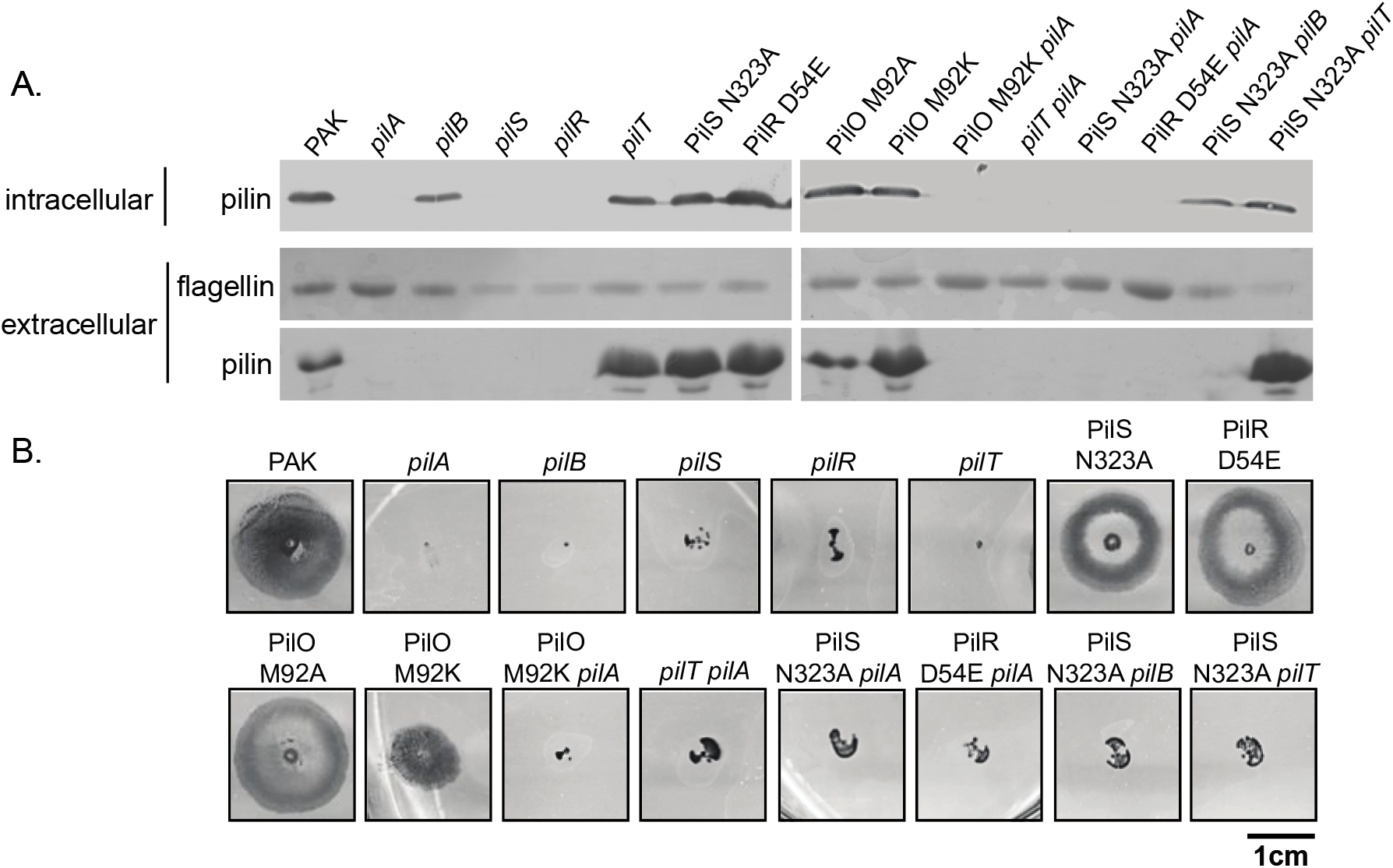
PilS N323A and PilR D54E point mutants are hyperpiliated, but have wild-type motility. **A.** The levels of intracellular (anti-PilA western blot) and extracellular pilins and flagellins (Coomassie-stained SDS-PAGE) of the mutants used in this study. The PilO M92K mutant was shown previously (41) to be hyperpiliated, while the isogenic M92A mutant resembles the WT. **B.** Representative twitching motility phenotypes of the mutants in panel A. The PilS N323A and PilR D54E mutants have levels of surface pili similar to a *pilT* mutant, but their twitching motility is similar to the WT. In addition to being hyperpiliated, the PilO M92K mutant has reduced twitching motility, consistent with a retraction defect (41).

### PilSR point mutants are less pathogenic towards C. elegans

To clarify the roles of piliation versus twitching motility in the pathogenicity of *P. aeruginosa*, we used a *C. elegans* slow killing infection model, where worms propagated on solid media feed on the bacterial strains of interest. As in previous studies (1, 42), non-piliated mutants including *pilA, pilS*, and *pilR* had slightly decreased pathogenicity compared to WT. In contrast, hyperactivation of PilS or PilR via the N323A or D54E point mutations, respectively, significantly decreased pathogenicity, with 50% killing that took on average 4 days longer than WT **(Figure 2A).** To confirm that this was not a strain-specific phenotype, we also tested the pathogenicity of PilS N323A and PilR D54E point mutants of the PA14 strain, which kills *C. elegans* more rapidly than PAK (43). We saw similar reductions in pathogenicity for the PA14 point mutants **(Supplementary Figure S1)**. These decreases in pathogenicity were not a result of altered growth of the point mutants on slow killing medium **(Supplementary Figure S2)**. PilA overexpression *in trans* in a WT background failed to cause hyperpiliation or reduce pathogenicity **(Supplementary Figure S3)**.

**Figure 2.**
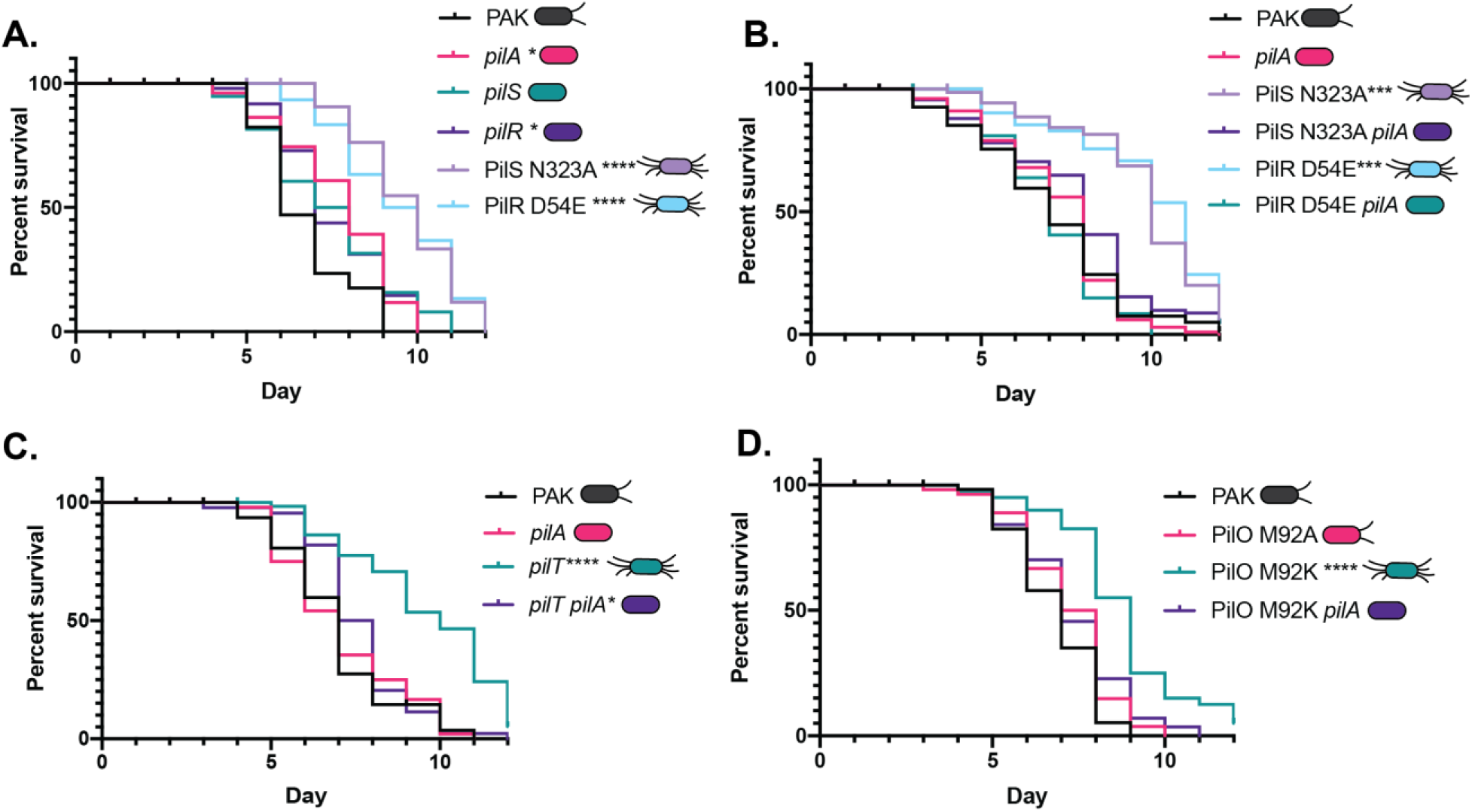
Hyperpiliation, not loss of pilus function, reduces pathogenicity in a pilin-dependent manner. **A.** The hyperpiliated PilS N323A and PilR D54E mutants are significantly less pathogenic in slow killing assays than WT PAK or its isogenic *pilA, pilS*, or *pilR* mutants; all of which lack surface pili. **B.** Deletion of *pilA* in the PilS N323A or PilR D54E backgrounds restores pathogenicity to levels similar to PAK and the *pilA* control. **C.** Loss of *pilA* in the *pilT* background, which is significantly less pathogenic than WT, restores pathogenicity to levels similar to WT and the *pilA* control. **D.** The hyperpiliated PilO M92K mutant is less pathogenic than PAK or an isogenic PilO M92A mutant, and pathogenicity is restored by deletion of *pilA* in the M92K background. * p<0.05, *** p<0.001, **** p<0.0001. Shown are representative datasets from triplicate or quadruplicate experiments; all replicates are available in the supplementary information **(Supplementary Figures S4-S7)**.

To clarify whether this phenotype was dependent on increased pilin expression, versus changes in expression of other members of the PilSR regulon (44), *pilA* was deleted in the PilS N323A and PilR D54E backgrounds. In both cases, the pathogenicity of the double mutants was comparable to that of PAK, suggesting that reduced pathogenicity in the PilSR point mutants was PilA-dependent **(Figure 2B)**.

### Other mutations that cause hyperpiliation similarly reduce pathogenicity

To understand how the level of surface piliation influences pathogenicity in *C. elegans*, we compared the pathogenicity of a *pilT* deletion mutant with a double mutant lacking both *pilT* and *pilA*. As expected, the *pilT* mutant was hyperpiliated and unable to twitch **(Figure 1BC).** In the slow killing assay, loss of *pilT* significantly reduced pathogenicity, while pathogenicity was restored to WT levels by deletion of both *pilA* and *pilT***(Figure 2C)**. As neither of these mutants can twitch, loss of pilus-mediated motility is not correlated with reduced pathogenicity in this infection model.

Each of the single mutations above has the capacity to either directly or indirectly modulate transcription of the pilin gene because of their effects on levels of pilin inventories in the inner membrane (9, 10, 45). For example, a *pilT* mutant has low levels of PilA in the inner membrane due to its inability to disassemble extended pili, while a *pilA* mutant has none. While both these mutations activate the PilSR regulon, the *pilT* and *pilA* mutants have different virulence phenotypes in *C. elegans*. To further separate the contributions to pathogenicity of changes in the levels of surface pili versus regulation of *pilA* transcription, we examined the pathogenicity of a strain that is hyperpiliated due to mutation in a gene outside of the known PilSR regulatory network (44). We previously identified two point mutations of residue M92 in the T4aP alignment subcomplex protein, PilO that differently affected surface piliation even though intracellular levels of PilA are similar (41). PilO M92A has no detectable impact on surface piliation or motility, while a charged residue at the same position, M92K, causes a hyperpiliated phenotype coupled with reduced motility **(Figure 1)**. Consistent with the results above, the hyperpiliated PilO M92K mutant, but not the M92A mutant, was significantly impaired in its ability to kill *C. elegans*, with the time to 50% mortality increased by an average of 2 days **(Figure 2D)**. Deletion of *pilA* in the PilO M92K background restored WT killing kinetics, further supporting our hypothesis that hyperpiliation is detrimental to pathogenicity in *C. elegans*.

### Overproduction of surface pili reduces swarming motility

We next tested whether swarming motility, a virulence-associated phenotype which is partially pilus-dependent in *P. aeruginosa* (46, 47), was affected by modulating the level of surface pili. We measured swarming motility over 48h and found that the non-piliated *pilA, pilS*, and *pilR* mutants all retained partial swarming motility, but the zones had a different morphology than those of WT, forming fewer but longer tendrils extending from the point of inoculation **(Figure 3)**. Conversely, functionally hyperpiliated PilS N323A and PilR D54E were both deficient in swarming motility, more closely resembling the non-flagellated *fliC* negative control. Loss of swarming motility in those strains was partially pilin dependent, as deletion of *pilA* in the PilS N323A or PilR D54E backgrounds restored swarming to levels similar to those of the *pilA* mutant **(Figure 3)**. Together, these data indicate that surface pili are required for WT swarming motility and that hyperpiliation is more detrimental to swarming than having no pili at all.

**Figure 3.**
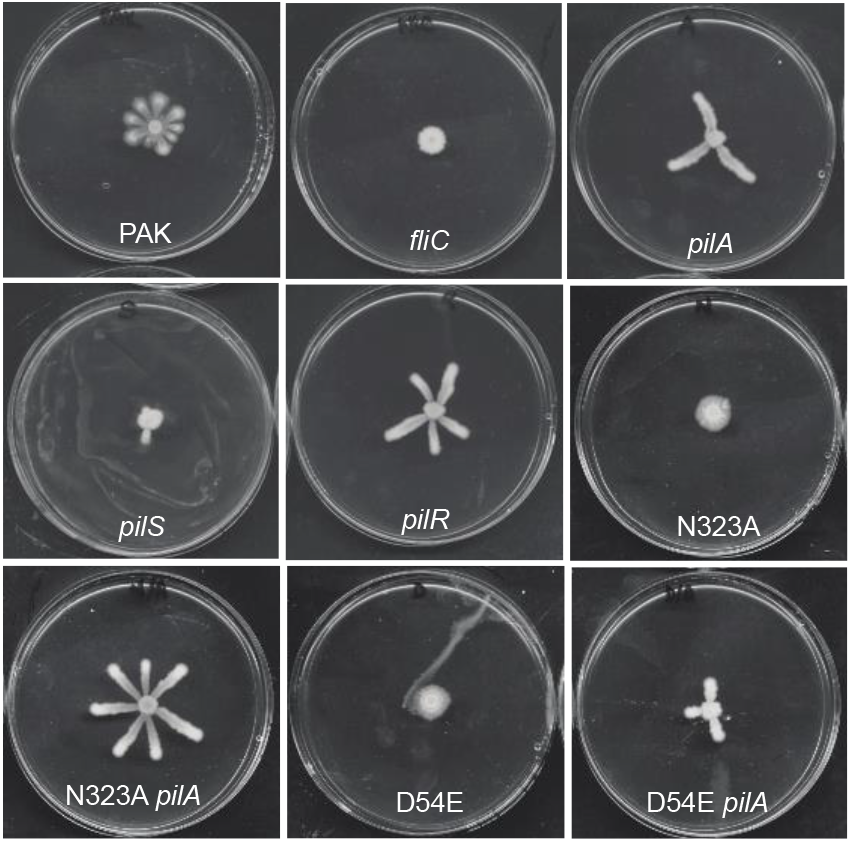
Hyperactivation of PilSR impairs swarming motility in a pilin-dependent manner. Swarming motility requires both flagella and pili. A flagellin mutant (*fliC*) is unable to swarm, while a *pilA* mutant has reduced swarming compared to WT PAK. Swarming of the *pilS* and *pilR* mutants, which lack pili, resembles that of the *pilA* mutant, with fewer tendrils. The hyperpiliated PilS N323A and PilR D54E mutants are unable to swarm, while deletion of *pilA* in those backgrounds leads to swarming patterns resembling the *pilA* mutant.

### Impaired Type III secretion contributes to the decreased pathogenicity of hyperpiliated mutants

To explain the decreased pathogenicity in hyperpiliated *P. aeruginosa* strains, we considered potential changes in the interaction between the bacteria and *C. elegans*. Efficient T3S relies on the establishment of intimate cell-cell contact between bacteria and host, and T3S is important for virulence in many infection models (36). The role of T3S in *P. aeruginosa* killing of nematodes has been controversial, and may depend on the specific *P. aeruginosa* strain tested. For example, studies using highly-virulent strain PA14 suggested that T3S has a negligible role in *C. elegans* slow killing (48), whereas others using the less-pathogenic PAO1 strain indicated that loss of T3S diminishes pathogenicity (49).

We tested the contribution of T3S to slow killing of *C. elegans* by strain PAK by generating a *pscN* deletion mutant lacking the T3S system ATPase, which prevents the secretion of toxic effectors as shown for *Yersinia enterocolitica* (50) and a number of other T3SS-expressing pathogens (51–54). Loss of *pscN* significantly reduced the pathogenicity of PAK in the slow killing assay **(Figure 4A).** In contrast, when the *pscN* mutation was introduced into the less pathogenic PilS N323A mutant, it did not further reduce pathogenicity. However, deleting *pscN* in a PilS N323A *pilA* double mutant – which has WT levels of virulence – decreased its pathogenicity. Together, these data suggest that T3S is important for *C. elegans* slow killing by PAK, and that hyperpiliation of *P. aeruginosa* may impair function of the T3SS.

**Figure 4.**
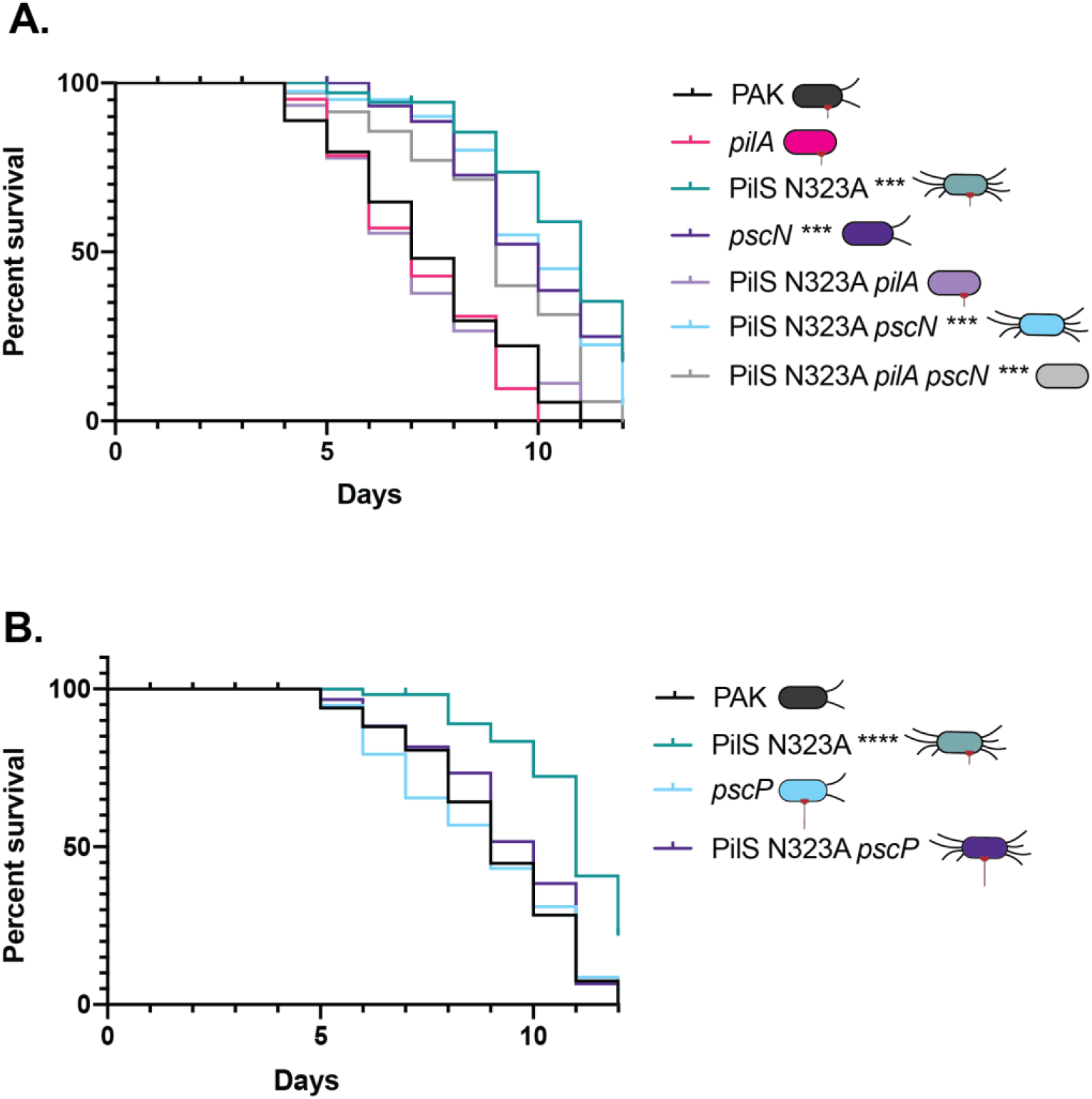
Increasing the length of T3SS needles restores pathogenicity in the hyperpiliated PilS N323A mutant. **A.** Deletion of *pscN*, encoding the T3SS ATPase, reduces pathogenicity of PAK towards *C. elegans*, showing that virulence of this strain is T3SS-dependent. Combining the *pscN* and PilS N323A mutations does not further decrease virulence. While loss of *pilA* in the N323A background increases pathogenicity, further deletion of *pscN* in the N323A *pilA* background reduces pathogenicity, confirming that virulence is T3SS-dependent. **B.** While deletion of *pscP*, encoding the T3SS ruler protein that controls needle length does not impair virulence of PAK towards *C. elegans*, deletion of this gene in the hyperpiliated N323A background increases pathogenicity. *** p<0.001, **** p<0.0001. These are representative datasets from triplicate experiments, and all replicates are available in the supplementary information **(Supplementary Figure S8)**.

We reasoned that if hyperpiliation of *P. aeruginosa* impeded the interaction of the T3SS system with the host, we may be able to overcome this problem if we could extend the length of T3SS needles. *pscP* encodes the T3S ruler protein, and mutants lacking PscP produce longer yet functional needle structures that can reach up to 1μm in length (55) – potentially sufficient to contact host cells even in hyperpiliated mutants. While deletion of *pscP* in a strain with WT piliation had no significant effect on pathogenicity, deleting *pscP* in the hyperpiliated PilS N323A strain restored pathogenicity to near-WT levels (**Figure 4B**). These data support the idea that having too many surface pili, even if they are functional in terms of retraction and twitching motility, interferes with the contact-dependent delivery of T3SS effectors, and thus, pathogenicity (**Figure 5**).

**Figure 5.**
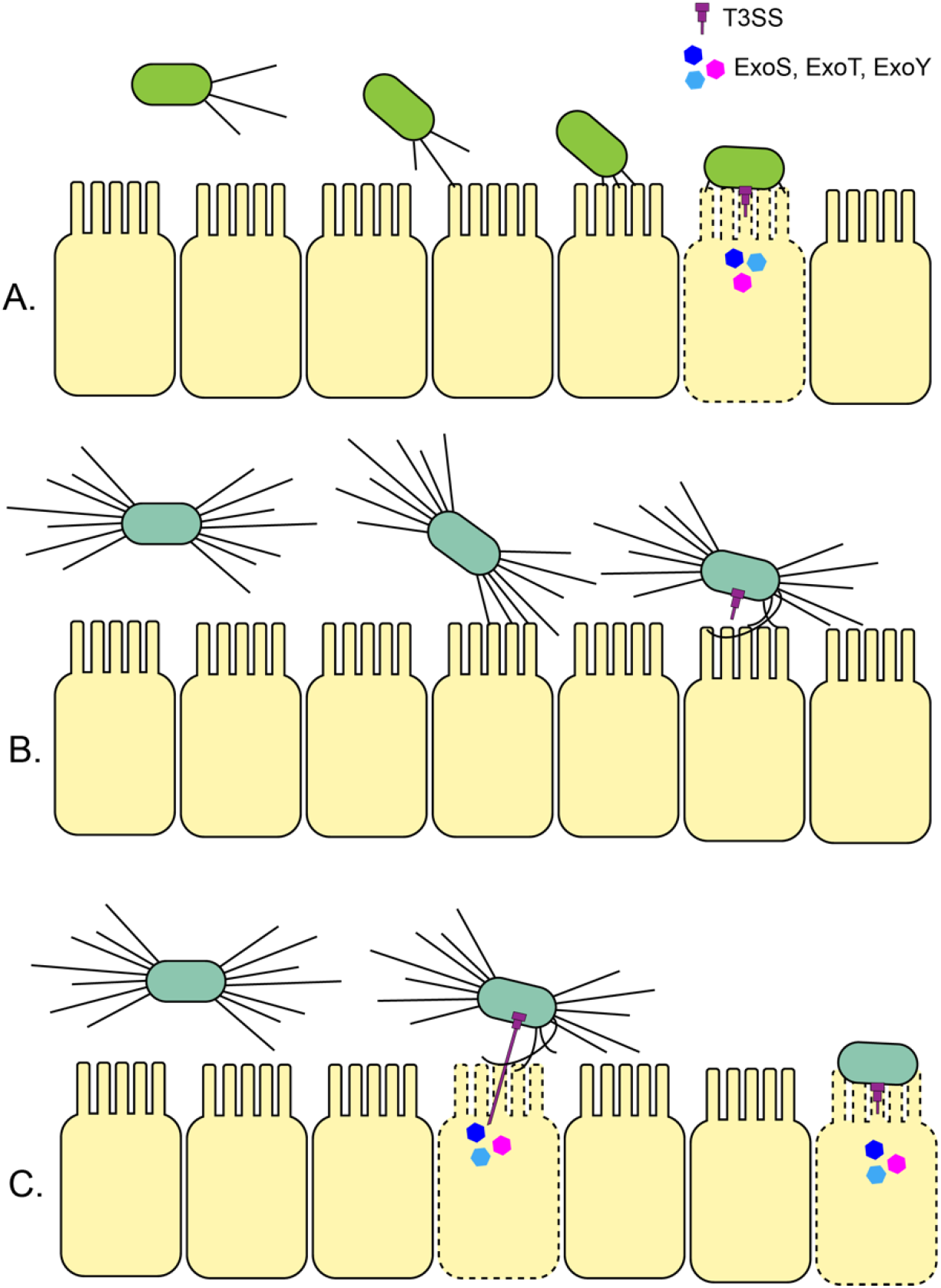
Hyperpiliation impedes engagement of the T3SS and reduces virulence. **A.** *P. aeruginosa* can retract its T4P upon contact with the epithelial cells lining the gut of the worms, leading to intimate engagement of the T3SS and injection of toxic effectors ExoS, ExoT and/or ExoY. **B.** The T3SS of hyperpiliated mutants (lacking the retraction ATPase PilT, or with specific point mutations in PilO, PilR, or PilS) are unable to effectively engage with host cells to inject effectors, and thus, are less pathogenic. **C.** In those hyperpiliated backgrounds, increasing T3SS needle length via deletion of the ruler protein gene *pscP*, or removing surface pili – which are not critical for pathogenicity in the slow killing model of *C. elegans* (56) – via *pilA* mutations restores virulence.

## DISCUSSION

T4aP are important virulence factors for *P. aeruginosa* and other bacterial pathogens (1, 2, 25). Prior studies of *pilT* mutants attributed their deficits in virulence to loss of pilus retraction and twitching motility, without considering the specific contribution of hyperpiliation (29, 30). Here, we showed that hyperpiliation, not loss of twitching motility or loss of *pilT* specifically, in *P. aeruginosa* substantially reduces its pathogenicity towards *C. elegans*. We propose that excess surface pili may allow for reversible attachment to the host, but impair the establishment of the intimate cell-cell interactions required for productive engagement of the contact-dependent T3SS.

Use of a set of hyperpiliated mutant strains with a range of twitching phenotypes enabled us to separate the contributions of surface piliation levels and motility to pathogenicity. Although *pilT*, PilS N323A, PilR D54E, and PilO M92K mutants are all hyperpiliated compared to WT, only *pilT* is completely deficient in twitching. PilS N323A and PilR D54E have fully functional pili and WT twitching motility, while PilO M92K has partial motility, suggesting a role for this protein – part of the alignment subcomplex that joins the pilus motor and secretin subassemblies – in modulating extension and retraction dynamics (41). Regardless of their twitching motility phenotypes, all hyperpiliated mutants had PilA-dependent defects in pathogenicity in *C. elegans*. These results help to explain the important role of PilSR in the regulation of *pilA* expression and modulation of the levels of surface piliation (11). If too many pili are produced, it can negatively impact multiple virulence phenotypes, including swarming motility and the ability to engage effectively with eukaryotic hosts.

Overexpression of PilA from an inducible plasmid in a WT background failed to cause hyperpiliation or reduce pathogenicity, likely because the normal PilSR regulatory circuit remained intact. When pilins are expressed *in trans*, they accumulate in the inner membrane and interact with PilS to decrease PilR activation, which balances pilin pools by reducing chromosomal expression of *pilA* (11). Using PilS N323A and PilS N323A *pilA* as representative hyperpiliated and non-piliated mutants, respectively, we showed there were no significant differences in growth rates or colonization capacity that would account for the decreased pathogenicity of PilS N323A and by extension, other hyperpiliated strains.

These data led us to conclude that excess surface pili can impair *P. aeruginosa’s* ability to kill *C. elegans* (**Figure 5**). All hyperpiliated mutants with reduced pathogenicity reverted to WT virulence when *pilA* was deleted. Notably, *pilT* and *pilA pilT* mutants, neither of which is capable of twitching, had divergent pathogenicity phenotypes. Those data indicated that – in contrast to previously published studies (22, 24, 28, 30) – loss of twitching motility is unlikely to be the cause of reduced infectivity in *C. elegans*. Rather, the increase in the amount of surface pili in *pilT* mutants is detrimental. Additional work is needed to determine if this holds true in other models of infection.

Interestingly, having too many pili had a more significant impact on pathogenicity in the *C. elegans* slow killing model than having no pili at all, although depending on the specific genetic lesion, some non-piliated mutants are more pathogenic than others (56). *P. aeruginosa* is actively ingested by the worms, rather than needing to establish contact with the host—a stage that normally relies on T4aP (1). Furthermore, the differences in pathogenicity between non-piliated and hyperpiliated strains – regardless of their ability to twitch – is obvious despite the proposed role of T4aP in the sensing of surface contact, upregulation of virulence, and surface display of the putative mechanosensory adhesin, PilY1 (7, 57). Recent work indicated that *pilT* is required for a c-di-GMP-dependent response to shear forces and surface sensing (58), suggesting that hyperpiliation may also disrupt signal cascades required for virulence in other contexts.

After demonstrating that hyperpiliation decreases pathogenicity in *C. elegans*, we ruled out reduced colonization as a possible mechanism. Prior studies showed that injection of the T3S effector, ExoS, requires T4P (33) and the retraction ATPase, PilT (34). Studies in PA14 indicated that while the T3SS is expressed during infection of *C. elegans*, it is not required for full pathogenicity (48), but also suggested that loss of the effector ExoU impaired PA14 virulence (59). In PAO1, the T3SS plays a major role, as loss of function significantly reduces pathogenicity (49). PAK is more closely related to PAO1 than to PA14, with PAK and PAO1 expressing the T3SS effectors ExoSTY, while PA14 expresses ExoSTU (60, 61). Consistent with the PAO1 data, a PAK *pscN* mutant had significantly reduced virulence towards *C. elegans*. Given the increased virulence of PA14 towards *C. elegans* relative to PAK and PAO1, it is possible that T3SS contributes to PA14 pathogenicity, but that more potent virulence factors produced by that strain kill *C. elegans* rapidly, before the contributions of T3SS effectors become obvious. Even so, our data show that hyperpiliated strains of PA14 are less pathogenic than WT.

The PilS N323A *pscN* double mutant had levels of pathogenicity similar to single PilS N323A and *pscN* mutants, and a PilS N323A *pscN pilA* triple mutant had decreased virulence compared to the PilS N323A *pilA* double mutant. Together, these data support a model where overproduction of surface pili, regardless of their functionality, impairs engagement of the T3SS, and therefore, pathogenicity in *C. elegans*. These data also refute previous conclusions that a functional T4P system is required for T3SS engagement (33, 34), as the PilS N323A *pilA* mutant lacks pili but was more pathogenic than the triple mutant that lacks PscN. Further supporting our hypothesis, deletion of *pscP* in the PilS N323A hyperpiliated background restored pathogenicity to near-WT levels, showing that the point mutant retains the capacity to be virulent as long as it is able to deploy the T3SS. PscP expression is independent of PilSR (44) and its deletion did not affect pathogenicity in an otherwise WT background.

While no clear connection between swarming motility and virulence has been made, swarming and T3S expression are positively correlated in laboratory and clinical strains (46, 62). Consistent with these data, PilS N323A and PilR D54E strains of PAK failed to swarm and were the least pathogenic of the strains tested. Deletion of *pilA* from those backgrounds restored swarming motility to that of a *pilA* single mutant, which swarms in a different pattern than WT, as pili play a supplemental role in PAK swarming motility (47).

Taken together, these data challenge the idea that loss of twitching motility or pilus retraction reduces *P. aeruginosa* pathogenicity in *C. elegans* and other eukaryotic models. Instead, we suggest that inappropriate increases in the amount of surface pili, even if they are functional, prevent effective engagement of the T3SS. With recent calls for the development of anti-virulence strategies to supplement the search for new antibiotics (63), these data demonstrate that dysregulating the surface expression of T4P could be a superior strategy for reducing pathogenicity than trying to block pilus biogenesis entirely.

## METHODS

### Strain construction and growth conditions

Bacterial strains and plasmids used in this study are summarized in **Supplementary Table S1.** Plasmid constructs were generated using standard cloning techniques and the restriction enzymes indicated in the primer table, **Supplementary Table S2,** and were introduced into either chemically competent *Escherichia coli* or *Pseudomonas aeruginosa* Strain K (PAK) using heat shock or electroporation, respectively. Deletion constructs were prepared by ligating fragments corresponding to 500 bp up- and downstream of the gene being deleted and cloning the resulting fusion into the pEX18Gm suicide vector. Deletion or point mutation constructs were introduced into *E. coli* SM10 and conjugated into the PAK parent strain as described previously (64). Mutations were confirmed using PCR and Sanger sequencing (Mobix, McMaster Genomics Facility, Hamilton). Unless otherwise indicated, strains were grown in Lennox broth (LB) (Bioshop) or on LB 1.5% agar plates. Where necessary, gentamicin was added to the media for selection at a concentration of 15μg/mL or 30μg/mL for *E. coli* and *P. aeruginosa*, respectively.

### Sheared surface protein preparations

Sheared surface protein preparations were performed as previously described (65). Briefly, strains of interest were streaked in a grid-like pattern on a 150mm x 15mm LB 1.5% agar plate and grown overnight at 37°C. Cells were scraped from the plate using a glass coverslip and resuspended in 4.5mL 1X phosphate-buffered saline (PBS; pH 7.4). Surface appendages were sheared from the 4.5mL of aliquoted suspension by vortexing for 30s and centrifuging at room temperature for 5 min at 16 000 x g. Supernatants were transferred to a new tube and centrifuged again for 20 min to remove any remaining cellular debris. Intracellular samples were prepared by diluting the remaining cell pellets to OD_600_=0.6 in 1X PBS and pelleting 1mL of cells. Pellets were resuspended and boiled in 100μL 1x SDS sample buffer (100μL of 1X SDS sample buffer (80mM Tris (pH 6.8), 5.3% (v/v) β-mercaptoethanol, 10% (v/v) glycerol, 0.02% (w/v) bromophenol blue and 2% (w/v) SDS). Sheared surface protein supernatants were transferred to a fresh tube and 5M NaCl and 30% polyethylene glycol (MW = 8000Da) were added to final concentrations of 0.5M and 3%, respectively. Samples were incubated on ice for 1.5 h, inverting occasionally to facilitate protein precipitation, then centrifuged at 16 000 x g for 30 min at 4°C to pellet sheared proteins. Supernatants were discarded and after allowing the protein pellets to air dry, they were resuspended and boiled in 150 μL 1x SDS sample buffer in preparation for SDS-PAGE analysis.

### Western Blotting

Samples were separated on 15% SDS PAGE and transferred to a nitrocellulose membrane. Following transfer, membranes were blocked in a 5% (w/v) skim milk powder in PBS. α-PAK PilA rabbit polyclonal antibodies (Cedarlane Laboratories, Burlington ON, Canada) and goat-α-rabbit secondary antibody conjugated to alkaline phosphatase (Sigma Aldrich, Oakville, ON, Canada) were used at 1:7500 and 1:5000 dilutions, respectively, and blots were developed according to manufacturer’s instructions. The data are representative of at least three independent experiments.

### Twitching motility assays

Twitching motility assays were performed as described previously (66). Briefly, a single colony of each strain of interest was stab inoculated to the agar-plastic interface of an LB 1% (w/v) agar plate. Plates were incubated at 37°C for 24-48h. Following incubation, the agar was carefully removed from the plate and discarded and twitching zones were visualized by staining the plastic plate with 1% (w/v) crystal violet. Plates were imaged using a standard scanner and twitching zones were measured using ImageJ (http://imagej.nih.gov/ij/, NIH, Bethesda, MD). At least 3 independent replicates were performed.

### Caenorhabditis elegans slow killing pathogenicity assays

Slow killing (SK) assays were performed as described previously (67). *Caenorhabditis elegans* strain N2 populations were propagated and maintained on Nematode Growth Media (NGM) plates inoculated with *E. coli* OP50. Eggs were harvested to obtain a synchronized population by washing worms and eggs from NGM plates with M9 (0.3% KH_2_PO_4_, 0.6% Na_2_PO_4_, 0.5% NaCl, 1mM MgSO_4_) buffer. Worms were degraded by adding buffered bleach, leaving only the eggs intact. Eggs were washed with M9 buffer and resuspended in M9 buffer with rocking overnight to hatch into L1 larvae. Synchronized L1 worms were plated on NGM plates for 45h at 20°C to develop into L4 worms. During this process, slow killing (SK) plates supplemented with 100μM 5-Fluoro-2’-deoxyuridine (FUDR) were inoculated with 100μL of a 5mL LB overnight culture of bacterial strains of interest and incubated at 37°C for 16-18h. Harvested and washed L4 worms (~30-40) were dropped by Pasteur pipette onto each SK plate. Using a dissecting microscope, plates were scored daily and dead worms were removed. Survival curves were prepared using Graphpad Prism 5.01 (La Jolla, CA) and statistically-significant differences in pathogenicity between strains were identified using Gehan-Breslow-Wilcoxon analysis.

### Swarming motility assays

Swarming motility assays were performed as in (47), with slight modifications. Briefly, 5mL liquid LB cultures were inoculated with *P. aeruginosa* strains of interest and grown overnight at 37°C with shaking. Swarming plates were prepared on the day the assays were to be performed, with 10x M8 salts (12.8% w/v Na_2_HPO_4_·7H_2_0, 3% KH_2_PO_4_, 0.5% NaCl, pH 7.4), diluted to a final 1x concentration with water, 0.5% agar, and autoclaved together. After autoclaving and cooling to ~60°C, media was supplemented with 2mM MgSO_4_, 0.2% glucose, 0.05% L-glutamic acid, and trace metals (composition available upon request). Media was pipetted into 100mmx15mm petri dishes in 20mL aliquots per plate and allowed to solidify upright at room temperature for 1.5 h. From each of the prepared overnight cultures, 3.5μL of inoculum was spotted in the centre of each plate, which were then incubated upright at 30°C in a humidity-controlled incubator for 48 h. Plates were imaged using a standard computer scanner and where required, surface area was calculated using ImageJ (http://imagej.nih.gov/ij/, NIH, Bethesda, MD). Representative images from four independent experiments are shown.

### Growth curves

Growth curves were performed by inoculating *P. aeruginosa* strains of interest in a 5mL overnight culture of liquid slow-killing assay media, omitting agar and FUDR. The following day, cultures were diluted 1:20 in fresh slow killing media (50μL culture in 950μL fresh media) and samples were plated in a 100 well honeycomb plate (Oy Growth Curves Ab Ltd) in 300 μL technical triplicates. Growth (OD_600_) was measured at 1 h intervals for 24 h using a Bioscreen C plate reader (Oy Growth Curves Ab Ltd) set with continuous shaking at 37°C, and curves were generated using GraphPad Prism 5.01 (La Jolla, CA). The mean and standard error of three independent biological replicates (nine total samples) are shown.

### C. elegans colonization assays

Colonization assays were performed using the methods described in (68). Briefly, worms and bacterial strains were prepared as for the Slow Killing assay. Synchronized L4 worms were seeded onto SK plates containing 30μg/mL gentamicin that had been preincubated overnight at 37°C with the *P. aeruginosa* strains of interest. For counter-selection against residual *E. coli*, all *P. aeruginosa* strains were transformed with the pBADGr plasmid, which confers gentamicin resistance. *C. elegans* was allowed to feed on the *P. aeruginosa* strains for five days. On each day, 10 worms were removed from the plate and washed three times in M9 buffer containing 1mM sodium azide to prevent expulsion of bacteria from the gut. A sample of the final wash was serially diluted and plated on LB 1.5% agar plates containing 30μg/mL gentamicin to estimate how many *P. aeruginosa* colony forming units (CFUs) remained on the exterior of the worms. Worms were then lysed by vortexing for 30 s in the presence of 4-6 stainless steel, 3.175 mm diameter beads (Lysing matrix S, MP Biomedicals, Canada). Lysates were serially diluted and plated on LB 1.5% agar with gentamicin to determine *P. aeruginosa* CFU/mL combined on the interior and exterior of the worms. These plates were grown overnight at 37°C and colonies were counted. To estimate *P. aeruginosa* CFU/mL inside *C. elegans*, the supernatant CFU/mL was subtracted from the lysate CFU/mL—which should contain internalized bacteria as well as remaining bacteria in the final wash. This difference, representing internalized bacteria, was plotted as CFU/mL over time. Measurements were made at 24 h intervals over 5 days. Experiment was repeated in triplicate and a one-way ANOVA statistical test with Dunnett’s post-test was used to assess significance at each time point, using PAK as the reference sample.

## ACKNOWLEDGEMENTS

We thank David Moskal for help with PilA overexpression studies, and Mercedes DiBernardo for assistance with worm experiments. This work was funded by a grant (PJT-156080) from the Canadian Institutes of Health Research to LLB. KJG held a Canadian Graduate Scholarship, SLNK held a Ontario Graduate Scholarship, and RPL held Michael G. DeGroote, CIHR, and Cystic Fibrosis Canada Fellowships.

